# Evidence for Similar Structural Brain Anomalies in Youth and Adult Attention-Deficit/Hyperactivity Disorder: A Machine Learning Analysis

**DOI:** 10.1101/546671

**Authors:** Yanli Zhang-James, Emily C Helminen, Jinru Liu, The ENIGMA-ADHD Working Group, Barbara Franke, Martine Hoogman, Stephen V Faraone

## Abstract

ADHD affects 5% of children world-wide. Of these, two-thirds continue to have impairing symptoms of ADHD into adulthood. Although a large literature implicates structural brain differences in the pathophysiology of the disorder, it is not clear if adults with ADHD have similar neuroanatomical impairments as those seen in children with recent reports from the large ENIGMA-ADHD consortium finding structural abnormalities for children but not for adults. This paper uses deep learning neural network classification models to determine if there are neuroanatomical changes in the brains of children with ADHD that are also observed for adult ADHD, and vice versa. We found that structural MRI data can significantly separate ADHD from control participants for both children and adults. Consistent with the prior reports from ENIGMA-ADHD, prediction performance and effect sizes were better for the child than the adult samples. The model trained on adult samples significantly predicted ADHD in the child sample, suggesting that our model learned anatomical features that common to ADHD in childhood and adulthood. These results support the continuity of ADHD’s pathophysiology from childhood to adulthood. In addition, our work demonstrates a novel use of neural network classification models to test hypotheses about developmental continuity.

## Introduction

ADHD is a common disorder affecting 5% of children and 3% of adults ^1^. It is associated with injuries ^2^, traffic accidents ^3^, increased health care utilization ^4, 5^, substance abuse ^6, 7^, criminality ^8^, unemployment ^9^, divorce ^10^, suicide ^11, 12^, AIDS risk behaviours ^13^ and premature mortality ^14^. The cost of adult ADHD to society is between $77.5 and $115.9 billion each year ^15^.

After decades of work documenting ADHD’s high heritability (76%) ^16^, we now know from a genome wide association study (GWAS) of over 50,000 samples that the common DNA variants associated with ADHD’s significant polygenic risk are highly enriched for genes expressed in brain ^17^ and that many of these genes are expressed in pathways crucial for neurodevelopment ^18^. A role for brain dysfunction in the aetiology of ADHD was suspected for some time by the mechanism of action of the medications that treat ADHD ^19^.

Because many structural magnetic resonance imaging (sMRI) studies had been published with conflicting results, the Enhancing Neuro Imaging Genetics Through Meta-Analysis (ENIGMA) ADHD Working Group create a large collaborative data set with sufficient power to detect small effects. The ENIGMA-ADHD working group found small, statistically significant sub-cortical volumetric reductions ^20^, cortical thinning and reduced surface area ^21^ to be associated with ADHD in children but not adults. Using data from the Allen Brain Atlas, Hess et al. ^22, 23^ subsequently compared gene expression profiles between brain regions that were and were not implicated in the ENIGMA-ADHD subcortical analyses. Gene expression profiles for three pathways were significantly correlated with ADHD-associated volumetric reductions: apoptosis, oxidative stress, and autophagy. These results suggest that variability of structural brain anomalies in ADHD might be explained, in part, by the differential vulnerability of these regions to mechanisms mediating apoptosis, oxidative stress, and autophagy. Hess et al.’s findings also provide some validation of the ENIGMA-ADHD findings.

An intriguing finding from the ENIGMA-ADHD results was that all significant findings were for childhood ADHD. They found no significant findings for adult ADHD. We use the term childhood ADHD to refer to ADHD ascertained in childhood, understanding from our meta-analysis that two-thirds will persist into young adulthood and that persistence continues to decline with age ^1^. We use the term adult ADHD to refer to childhood onset ADHD that has persisted into adulthood, which is how it is defined in DSM 5 and in the ENIGMA-ADHD sub-studies. The ENIGMA-ADHD results are partly consistent with longitudinal studies show decreases in ADHD vs. control differences in regional volumes and cortical thicknesses ^24–26^. Those ADHD participants whose brains become more neurotypical were more likely than others to show remission of symptoms ^27, 28^. But, although these longitudinal studies show reductions in case vs control differences, they also suggest that those difference should be evident to some degree in cases that persist into adulthood.

Although the expectation of finding substantial continuity between childhood and adult ADHD has been widely accepted ^29–31^ and recently confirmed by a large GWAS ^32^, this idea has been challenged ^33^. Thus, given these prior data and the controversy about the continuity of ADHD into adulthood, we sought to test the idea that the ADHD-associated volumetric reductions seen in children with ADHD would be detected in adults with ADHD by applying machine learning algorithms. Given that symptoms and impairments persist into adulthood for most children with ADHD ^1, 34^, we hypothesized that ADHD-related brain structure differences in adults would be consistent with those observed in children.

## Materials and Methods

### MRI Samples

The current study was approved by all contributing members of the ENIGMA-ADHD Working Group, which provided T1-weighted structural MRI (sMRI) data from 4,183 subjects from 35 participating sites (by Aug 2019). Each participating site had approval from its local ethics committee to perform the study and to share de-identified, anonymized individual data. Images were processed using the consortium’s standard segmentation algorithms in FreeSurfer (V5.1 and V5.3) ^20^. 151 variables were used including 34 cortical surface area, 34 cortical thickness measurements and 7 subcortical regions from each hemisphere, and intracranial volume (ICV). Subjects missing more than 50% of variables were removed. Remaining missing values and outliers (outside of 1.5 times the interquartile range (iqr 1.5)) were replaced with imputed values using multiple imputation with chained equations in STATA15. The final ML dataset consisted 4,042 subjects from 35 sites, among which 45.8% were non-ADHD controls (n=1,850, male to female ratio (m/f) = 1.42) and 54.2% ADHD participants (n=2,192, m/f=2.79). Ages ranged from four to 63 years old; 60.7% were children (age<18 years, n=2,454) and 39.3% were adults (age≥18 years, n= 1,588). ADHD diagnosis was significantly biased by sex (X^2^_(1)_ = 66.9, p<.0001), sites (X^2^_(1)_ = 146.73, p<.0001), age (X^2^_(1)_ = 4.28, p=0.04).

To balance the confounding factors, we took the following steps. First, we randomly assigned samples to training (~70%), validation (~15%), and test (~15%) subsets within each diagnosis, sex, age subgroup (child vs adult) and site to ensure that the train/validation/test subsets have the same composition of these variables. 12 sites that provided only cases or only controls (total 203 subjects) were excluded during the initial train/validation/test split because their samples cannot provide an unbiased learning during the training and validation steps. These samples were added to the test set for final test evaluation. Supplementary Table 1 shows the sample splitting from each site. Next, we balanced the training set for the case and control groups within each sex, age and site subgroup by random oversampling of the under-represented diagnostic group, a procedure commonly used to deal with class imbalance. The resulting balanced training set is described in Table 1. The validation and test sets were not balanced by age, sex and site, however due to our sample splitting procedures, they contain the same demographic samples as the training set. In addition, the test set also contains samples from sites that had been excluded from the training set due to not having a site-specific control group.

**Table 1.** Training set sample characteristics after balancing for age and sex.

| Diagnosis |  | Child (Age <18) |  | Adult (Age ≥18) |  |
| --- | --- | --- | --- | --- | --- |
|  |  | Female | Male | Female | Male |
| Control | N of Subjects | 352 | 714 | 224 | 373 |
|  | Mean Age | 11.3 | 11.6 | 31.9 | 28.1 |
|  | SD of Age | 2.9 | 2.9 | 11.5 | 9.4 |
| ADHD | N of Subjects | 352 | 714 | 224 | 373 |
|  | Mean Age | 11.0 | 11.8 | 32.2 | 28.8 |
|  | SD of Age | 2.6 | 2.7 | 10.6 | 9.4 |

\*Note: SD, standard deviation; N, total numbers.

### Feature preprocessing

Principal factors factor analysis (PFFA) with varimax rotation on sMRI features on the training set identified 46 factors that explained >90% of the variance. Factor scores were computed for all subjects based on the training set PFFA. All input features were scaled based on the training set’s minimum and maximum values.

### Neural Network Framework

We implemented multilayer perceptron (MLP) neural network models using the Keras API (version 2.3.1) and the TensorFlow library (version 1.14.0). We used HyperOpt ^35^ to search the neural network hyperparameter spaces including numbers of layers, numbers of units and dropout rates in each layer, learning rate and batch normalization size. We also tested different activation functions and optimizers. We used binary cross entropy as the loss function. Early stopping was implemented to avoid overfitting. Best model architecture and hyperparameters were chosen based on the lowest total validation loss. Final test scores were obtained on the test set with ensemble learning approach ^36^. All machine learning algorithms were written in Python 3.5.

### Analysis Pipeline

Our analysis pipeline starts with two base models that used data from the corresponding age groups during the model training and validation phase and tested also on data from their corresponding age groups. The child model used only child samples during model training, validation and hyperparameter optimization, and tested on child test set. The adult model, similarly, was trained and validated on the adult samples and tested on the adult test set. We examined models using MRI features only, as well as those included age and sex information.

Next, we tested if the model trained and validated on the adult samples, the adult model, could be used to predict child ADHD, and vice versa. We hypothesized that if the ADHD vs. control sMRI differences seen in children are also present in adult ADHD brains, then the base models for each age group should be able to predict ADHD in the other age group. To create the largest test sets possible, we tested the child model on all the adult samples, and the adult model on all the child samples.

### Model evaluation

The sigmoid function in the output layer of the neural network generates a continuous score that assesses the probability for each individual to be classified as ADHD. We name this continuous output the brain risk score. Using the brain risk scores, we calculated Cohen’s *d* effect sizes for child and adult test sets. We computed receiver operating characteristic (ROC) curves and used the area under the ROC curve (AUC) as our primary measure of accuracy. The AUC and its confidence intervals were calculated in Stata 15 using the empirical method and compared with nonparametric approach by DeLong et al. ^37^. We also computed precision-recall (PR) curves and reported the area under the PR curves, as well as the Brier loss for the final models as measures of accuracy and goodness of fit.

## Results

Figure 1A Top portion shows the test set AUCs (as dots) and their 95% confidence intervals (as horizontal lines) for the base models using only MRI factors. The model trained and validated on child data predicted child ADHD with a significant AUC 0.64 (95%CI 0.58-0.70). In contrast, the model trained and validated on adult data resulted in a marginally significant AUC (0.56, 95%CI 0.49− 0.62, p= 0.057). The difference between the two base models’ AUCs was also marginally significant (X^2^_(1)_ = 3.4, p= 0.065). The areas under the precision-recall curve (AUPRC) were higher for the adult model (AUPRC = 0.74) than the child model (AUPRC= 0.68). Using the model predicted brain risk scores, we calculated the Cohen’s *d* effect sizes in the test set to be 0.47 for child samples (95%CI: 0.27 - 0.68) and 0.15 (−0.08 - 0.39) for the adult samples.

**Figure 1.**
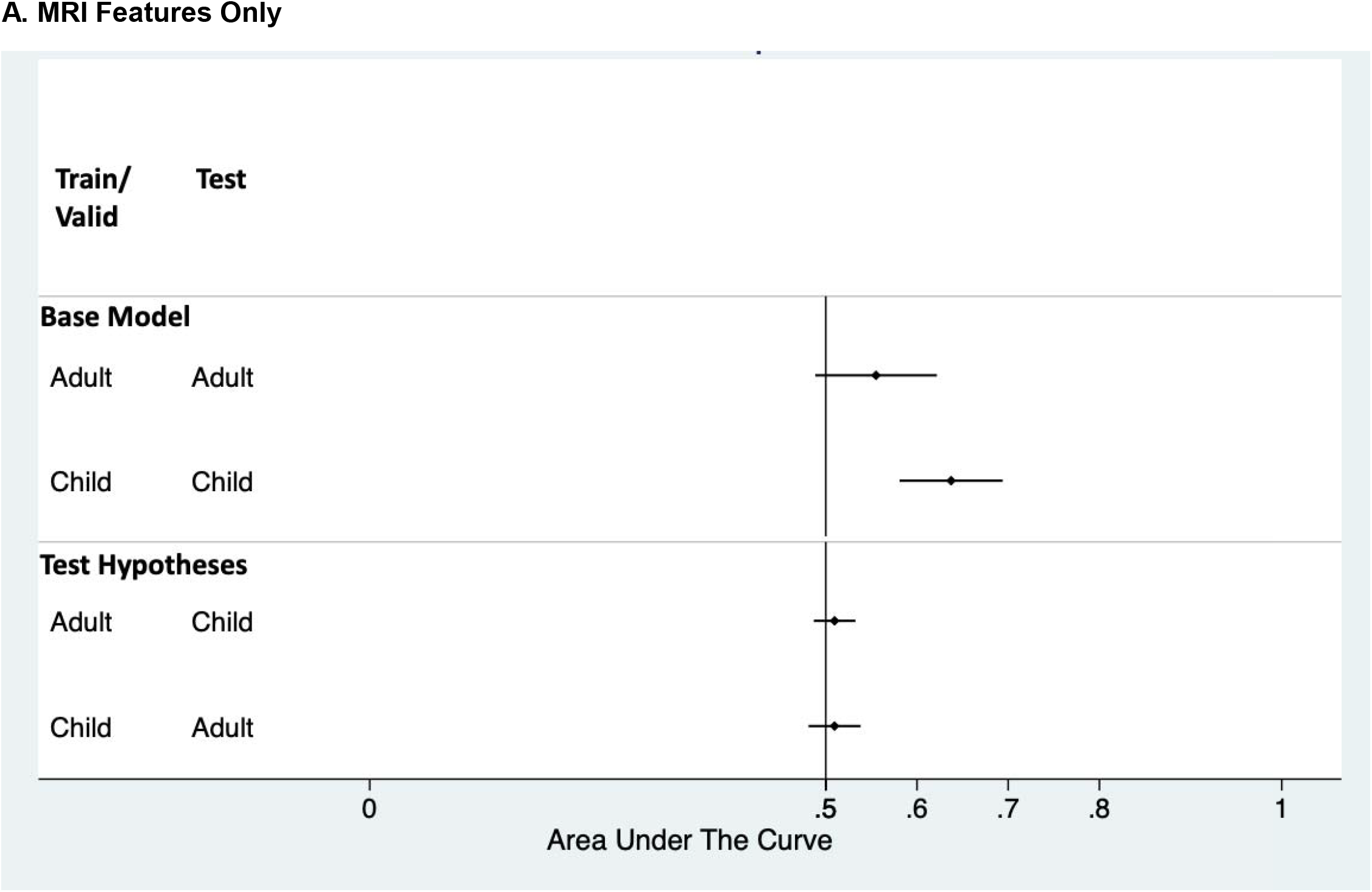
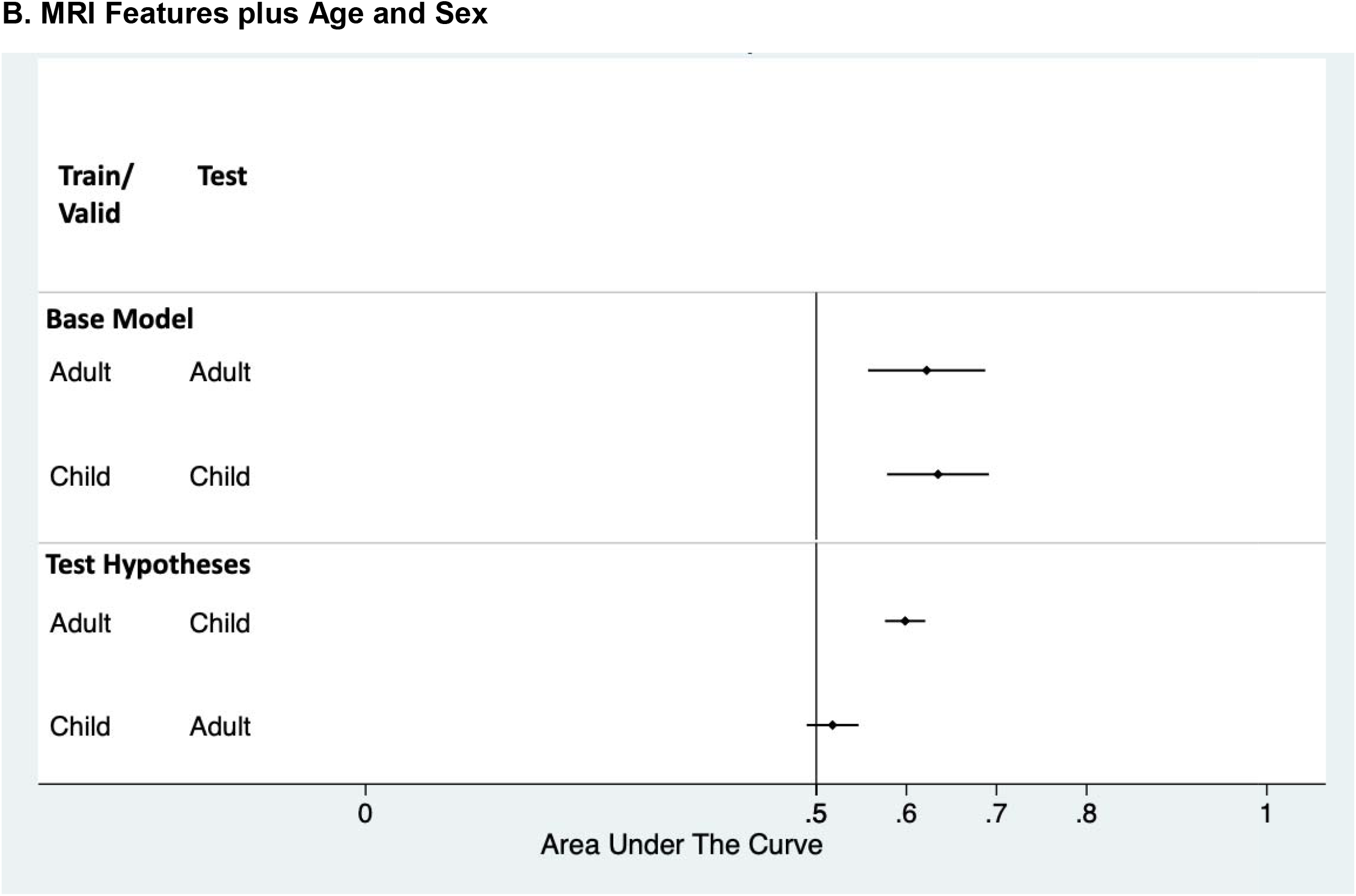
Area Under the Receiver Operating Characteristic Curve for the Test Results. Area under the receiver operating characteristic curve (AUC) accuracy statistics for the held-out test results were plotted (as dots) with their 95% confidence intervals (as horizontal lines). The vertical line at an AUC of 0.5 indicates a chance level of diagnostic accuracy. If the 95%CI does not overlap with the 0.5 vertical line, it indicates significant predictive accuracy. **A.** AUC comparison of the models using only MRI features. **B.** AUC comparison of the models using MRI features plus age and sex. In both **A** and **B**, **Top** portion shows the base models, where models were trained and validated in child or adult samples and tested on their corresponding age groups; **Bottom** portion tests the hypotheses that if model trained/validated on child samples can also predict adult ADHD and vice versa. Note that test sample consists of combined training, validation and test sets from the other age group because they are not used in the model optimization and training.

After adding age and sex as predictors, the adult model (Figure 1B Top) increased the AUC to 0.62 (95%CI 0.56 - 0.69). The AUPRC for the adult model also slightly increased (to 0.79. Adding age and sex as predictors to the child model did not affect either the AUC, nor the AUPRC. The Cohen’s *d* effect sizes in the test set were 0.48 for children (95%CI: 0.27-0.69) and 0.39 (0.15-0.63) adults. All above models had similarly small Brier scores (0.25).

It is worth noting that, because the training samples had been balanced for age and sex, these variables are not predictive of ADHD for either the child and adult test sets. To verify this, linear regression using only age and sex and their interactions to predict ADHD in the child and adult samples resulted in non-significant AUCs (child AUC 0.51, 95%CI: 0.45-0.57; adult AUC 0.46, 95%CI: 0.39-0.53).

### Tests of Hypotheses

For models using only MRI features, neither the adult, nor child models were successful at predicting ADHD in the other age group (Figure 1A Bottom). However, the adult model that used both MRI features and age and sex was able to predict the child samples significantly (AUC = 0.60, 95%CI: 0.58-0.62, Figure 1B Bottom). In contrast, the child model did not significantly predict ADHD when applied to the adult samples (AUC = 0.52, 95%CI: 0.49, 0.55, Figure 1B Bottom).

## Discussion

Consistent with previous ENIGMA ADHD findings ^20, 38^, we found that the ability of structural MRI data to discriminate people with and without ADHD is much stronger for children than adults, which is consistent with a broader literature showing that ADHD-associated structural brain differences diminish with age ^24–28^. While the ENIGMA ADHD study did not find any significant differences between ADHD and control subjects for adults, our adult model did achieve a significant AUC 0.62 (95%CI 0.56−0.69) and a high area under the PR curve (AUPRC=0.79). Consistent with the ENIGMA findings, our model-predicted brain risk scores had a larger effect size for the children than adults in both the models using MRI features and those with age and sex added. Notably, our effect sizes were two times greater than the largest of those individual regions reported in prior ENIGMA ADHD studies for both children (Cohen’s *d* = −0.21) and adults (Cohen’s *d* =−.16)^20, 38^.

Although the results from the child and adult base models indicate that sMRI data are not sufficiently predictive to be useful in clinical practice, they provide three crucial pieces of evidence that will be useful in future attempts at predictive modeling. First, our results are the first to confirm in the largest possible adult ADHD MRI sample available, that adults with ADHD differ significantly from adults without ADHD on sMRI features.

Second, the improvements we found by adding age and sex to the adult model indicate that these demographic variables must moderate the predictive ability of sMRI features. We believe that these demographics moderate the sMRI effects because our regression models show that the demographic variables on their own have no predictive utility (which was fixed in advance by balancing the case and control training samples by age and sex). Third, we have shown that machine learning methods dramatically increase the ADHD vs. Control effect size compared with the univariate ENIGMA analyses.

The results from our hypothesis testing provide further information that is useful in understanding the continuity of child and adult ADHD. Consistent with our hypothesis, the adult model, trained only on adult samples, significantly predicted ADHD in the child samples. This suggests that the adult model was able to learn combinations of structural features that are relevant for discriminating the structural MRI scans from children with and without ADHD. This implies that some of ADHD’s sMRI pathophysiology that is relevant for persistent cases is also relevant in childhood (only some of which will be persistent into adulthood). This conclusion must, however, be considered equivocal because the child model did not successfully predict ADHD in the adult samples. To resolve this issue, future studies will need to find a way to better discriminate sMRI features associated with the onset of ADHD and those associated with the persistence of ADHD.

Our work should be interpreted in the context of several limitations. First, because we combined data across many sites, we inherit all the limitations of the original studies. Heterogeneity of methods across studies may have added noise to the combined data set that made it difficult to discriminate the data from people with and without ADHD. Second, we only used structural imaging data. Incorporating other imaging modalities might provide clearer results and conclusions. Third, we used pre-defined structures from ENIGMA standard image processing pipeline as features. It is possible that other methods such as one using 3D images as input features, in a convolutional neural network would uncover useful features leading to increased classification accuracy. However, the 3D images are not available. Finally, our use of neural networks makes it difficult to clarify the importance of each brain region in the model’s algorithm.

Despite these limitations, we have shown that a neural network approach is able to detect case-control sMIR differences in adults with ADHD that could not be detected with standard analyses. We have also provided some evidence for the continuity of sMRI findings from childhood into adulthood.

## Funding and Disclosures

Dr. Faraone is supported by the European Union’s Seventh Framework Programme for research, technological development and demonstration under grant agreement no 602805, the European Union’s Horizon 2020 research and innovation programme under grant agreements No 667302 & 728018 and NIMH grants 5R01MH101519 and U01 MH109536-01. Dr. Franke is supported by a personal Vici grant (016-130-669) and Dr. Hoogman from a personal Veni grant (91619115), both from the Netherlands Organization for Scientific Research (NWO). The ENIGMA Working Group gratefully acknowledges support from the NIH Big Data to Knowledge (BD2K) award (U54 EB020403 to Paul Thompson).

Dr. Barbara Franke has received educational speaking fees from Shire and Medice.

Dr. Stephen V Faraone received income, potential income, travel expenses continuing education support and/or research support from Takeda, OnDosis, Tris, Otsuka, Arbor, Ironshore, Rhodes, Akili Interactive Labs, Enzymotec, Sunovion, Supernus and Genomind. With his institution, he has US patent US20130217707 A1 for the use of sodium-hydrogen exchange inhibitors in the treatment of ADHD. He also receives royalties from books published by Guilford Press: Straight Talk about Your Child’s Mental Health, Oxford University Press: Schizophrenia: The Facts and Elsevier: ADHD: Non-Pharmacologic Interventions. He is Program Director of www.adhdinadults.com.

Dr. Asherson has served as a consultant and as a speaker at sponsored events for Eli Lilly, Novartis, and Shire, and he has received educational/research awards from Eli Lilly, GW Pharma, Novartis, QbTech, Shire, and Vifor Pharma.

Dr. Banaschewski has served in an advisory or consultancy role for Actelion, Eli Lilly, Hexal Pharma, Lundbeck, Medice, Neurim Pharmaceuticals, Novartis, Oberberg GmbH, and Shire; he has received conference support or speaking fees from Eli Lilly, Medice, Novartis, and Shire; he has been involved in clinical trials conducted by Shire and Viforpharma; and he has received royalties from CIP Medien, Hogrefe, Kohlhammer, and Oxford University Press.

Dr. Bellgrove has received speaking fees and travel support from Shire.

Dr. Biederman has received research support from AACAP, Alcobra, the Feinstein Institute for Medical Research, the Forest Research Institute, Genentech, Headspace, Ironshore, Lundbeck AS, Magceutics, Merck, Neurocentria, NIDA, NIH, PamLab, Pfizer, Roche TCRC, Shire, SPRITES, Sunovion, the U.S. Department of Defense, the U.S. Food and Drug Administration, and Vaya Pharma/Enzymotec; he has served as a consultant or on scientific advisory boards for Aevi Genomics, Akili, Alcobra, Arbor Pharmaceuticals, Guidepoint, Ironshore, Jazz Pharma, Medgenics, Piper Jaffray, and Shire; he has received honoraria from Alcobra, the American Professional Society of ADHD and Related Disorders, and the MGH Psychiatry Academy for tuition-funded CME courses; he has a financial interest in Avekshan, a company that develops treatments for ADHD; he has a U.S. patent application pending (Provisional Number #61/233,686) through MGH corporate licensing, on a method to prevent stimulant abuse; and his program has received royalties from a copyrighted rating scale used for ADHD diagnoses, paid to the Department of Psychiatry at Massachusetts General Hospital by Ingenix, Prophase, Shire, Bracket Global, Sunovion, and Theravance.

Dr. Brandeis has served as an unpaid scientific consultant for an EU-funded neurofeedback trial.

Dr. Buitelaar has served as a consultant, advisory board member, and/or speaker for Eli Lilly, Janssen-Cilag, Medice, Roche, Shire, and Servier.

Dr. Coghill has served in an advisory or consultancy role for Eli Lilly, Medice, Novartis, Oxford Outcomes, Shire, and Viforpharma; he has received conference support or speaking fees from Eli Lilly, Janssen McNeil, Medice, Novartis, Shire, and Sunovion; and he has been involved in clinical trials conducted by Eli Lilly and Shire.

Dr. Dale is a founder of and holds equity in CorTechs Labs, Inc., and has served on the scientific advisory boards of CorTechs Labs and Human Longevity, Inc., and he receives funding through research grants with GE Healthcare.

Mr. Earl is co-inventor of the Oregon Health and Science University Technology #2198 (co-owned with Washington University in St. Louis), FIRMM: Real time monitoring and prediction of motion in MRI scans, exclusively licensed to Nous, Inc.) and any related research. Any potential conflict of interest has been reviewed and managed by OHSU.

Dr. Fair is a founder of Nous Imaging, Inc.; any potential conflicts of interest are being reviewed and managed by OHSU.

Dr. Haavik has received speaking fees from Biocodex, Eli Lilly, HB Pharma, Janssen-Cilag, Medice, Novartis, and Shire.

Dr. Hoekstra has received a research grant from and served on the advisory board for Shire.

Dr. Karkashadze has received payment for article authorship and speaking fees from Sanofi and from Pikfarma.

Dr. Konrad has received speaking fees from Eli Lilly, Medice, and Shire.

Dr. Kuntsi has received speaking honoraria and advisory panel payments for participation at educational events sponsored by Medice; all funds are received by King’s College London and used for studies of ADHD.

Dr. Lesch has served as a speaker for Eli Lilly and has received research support from Medice and travel support from Shire.

Dr. Mattos has served on speakers’ bureau and/or as a consultant for Janssen-Cilag, Novartis, and Shire and has received travel awards from those companies to participate in scientific meetings; the ADHD outpatient program (Grupo de Estudos do Déficit de Atenção/Institute of Psychiatry) chaired by Dr. Mattos also received research support from Novartis and Shire.

Dr. Mehta has received research funding from Lundbeck, Shire, and Takeda and has served on advisory boards for Lundbeck and Autifony.

Dr. Ramos-Quiroga has served on the speakers bureaus and/or as a consultant for Almirall, Braingaze, Eli Lilly, Janssen-Cilag, Lundbeck, Medice, Novartis, Shire, Sincrolab, and Rubió; he has received travel awards for taking part in psychiatric meetings from Eli Lilly, Janssen-Cilag, Medice, Rubió, and Shire; and the Department of Psychiatry chaired by him has received unrestricted educational and research support from Actelion, Eli Lilly, Ferrer, Janssen-Cilag, Lundbeck, Oryzon, Psious, Roche, Rubió, and Shire.

Dr. Reif has received honoraria for serving as speaking or on advisory boards for Janssen, Medice, Neuraxpharm, Servier and Shire.

Dr. Rubia has received speaking fees form Shire and Medice and a grant from Eli Lilly.

Dr. Thompson has received funding support from Biogen.

Dr. Van Erp has served as consultant for Roche Pharmaceuticals and has a contract with Otsuka Pharmaceutical, Ltd.

Dr. Walitza has received lecture honoraria from Eli Lilly and Opopharma, support from the Hartmann Müller, Olga Mayenfisch, and Gertrud Thalmann foundations, and royalties from Beltz, Hogrefe, Kohlhammer, Springer, and Thieme.

Dr. Yanli Zhang-James is supported by the European Union’s Seventh Framework Programme for research, technological development and demonstration under grant agreement no 602805 and the European Union’s Horizon 2020 research and innovation programme under grant agreements No 667302.

Emily C Helminen, Jinru Liu, Dr. Martine Hoogman and other contributing members of the ENIGMA-ADHD Working Group declare no conflict of interest.

## Acknowledgements

We thank Margaret Mariano and Patricia Forken for administrative assistance and proofreading the manuscript.

---

**Supplementary Table 1.** Total sample and train/validation/test splits from each site.

|  |  |  |  |  |  |
| --- | --- | --- | --- | --- | --- |
| **Sites** | **Training** | **Validation** | **Test** | **Excluded** | **Total** |
| ACPU | 46 | 11 | 9 | 1 | 67 |
| ADHD_WUE | 75 | 18 | 14 | 0 | 107 |
| ADHD-DUB1 | 56 | 14 | 10 | 0 | 80 |
| ADHD-DUB2 | 0 | 0 | 0 | 20 | 20 |
| ADHD-Mattos | 0 | 0 | 0 | 31 | 31 |
| ADHD200_KKI | 61 | 14 | 10 | 0 | 85 |
| ADHD200_NYU | 158 | 36 | 31 | 3 | 228 |
| ADHD200_OHSU | 61 | 16 | 12 | 0 | 89 |
| ADHD200_Peking | 139 | 31 | 27 | 0 | 197 |
| ADHDKonrad | 100 | 24 | 20 | 1 | 145 |
| ADHD_Rubia1 | 45 | 11 | 9 | 6 | 71 |
| ADHD_Russia | 0 | 0 | 0 | 10 | 10 |
| Barcelona | 51 | 12 | 10 | 0 | 73 |
| Bergen_SVG | 35 | 10 | 6 | 0 | 51 |
| Bergen_adultADHD | 55 | 15 | 11 | 0 | 81 |
| CAPS_UZH | 41 | 9 | 5 | 0 | 55 |
| DAT_london | 38 | 11 | 7 | 0 | 56 |
| Dundee | 32 | 8 | 4 | 1 | 45 |
| EPOD | 0 | 0 | 0 | 92 | 92 |
| Hartford_Olin | 125 | 31 | 25 | 0 | 181 |
| IMpACT_NL | 188 | 42 | 38 | 0 | 268 |
| MGH_ADHD | 100 | 24 | 20 | 0 | 144 |
| MTA | 91 | 21 | 17 | 0 | 129 |
| NICAP | 102 | 24 | 20 | 0 | 146 |
| NIH | 282 | 63 | 59 | 9 | 413 |
| NYU_ADHD | 56 | 14 | 10 | 0 | 80 |
| NeuroImage_ADAM | 118 | 29 | 21 | 0 | 168 |
| NeuroImage_NIJM | 120 | 30 | 22 | 0 | 172 |
| OHSU2018 | 161 | 36 | 32 | 0 | 229 |
| SAOPAULO | 92 | 22 | 18 | 1 | 133 |
| Sussex | 40 | 11 | 7 | 0 | 58 |
| Tuebingen | 0 | 0 | 0 | 28 | 28 |
| UAB-ADHD | 138 | 34 | 26 | 0 | 198 |
| UCHZ | 54 | 16 | 8 | 0 | 78 |
| ZI-CAPS | 24 | 7 | 3 | 0 | 34 |
| **Total** | **2,684** | **644** | **511** | **203** | **4,042** |

## Notes

#### Summary of Updates

Revised analysis and results

